# Kinetics of zoospores approaching a root using a microfluidic device

**DOI:** 10.1101/2023.06.21.545863

**Authors:** C Cohen, FX Gauci, X Noblin, E Galiana, A Attard, P Thomen

## Abstract

*Phytophthora* species are plant pathogens that cause considerable damage to agrosystems and ecosystems, and have a major impact on the economy. Infection occurs when their biflagellate zoospores move and reach a root on which they aggregate. However, the communication between the plant and the zoospores and how this communication modifies the behavior of the swimming zoospores is not yet well characterized. Here we show that using a microfluidic device comprising a growing *Arabidopsis thaliana* root, we are able to study the kinetics of *Phytophthora parasitica* zoospores approaching the root and aggregating on a specific area, in real time. We show that the kinetics of zoospores is modified only below a distance of about 300 *μ*m from the center of aggregation, with a decrease in the speed coupled with an increase in the number of turns made. In addition, we show that the rate of aggregation is constant throughout the experiment, approximately one hour, and depends on the density of zoospores. The rate is consistent with a random encounter of zoospores with the root, indicating that no long range signal is evidenced in our set-up.

## Introduction

The genus *Phytophthora* groups filamentous eukaryotic microorganisms belonging to the class Oomycetes. A number of *Phytophthora* species are plant pathogens that cause considerable damage to agrosystems and ecosystems: diseases caused by *Phytophthora* therefore have a major impact on the economy and constitute a threat to food security worldwide [12, 9].

In most cases, dispersal and primary infection are mediated by airborne sporangia or waterborne zoospores [22, 38]. Zoospores are biflagellate spores that spread in water. They are capable of reaching a velocity of 250 *μ*m/s [1] through thin films of water, water droplets on leaves, or through the pores of moist soils. When the zoospores reach plant roots, they stop swimming and release their flagella to produce a primary cell wall and become germinating cysts that are able to penetrate host tissue [28, 34]. Then they begin hyphal growth within the infected plant.

In the last decade, the main knowledge concerning plant-pathogen interactions has focused on the penetration and the colonization steps. It shows that plants and microorganisms establish communication throughout the infection, and that this dialogue is crucial for the outcome of the disease [21]. Plant roots secrete a variety of exudates in the rhizosphere that can be perceived by pathogens as signaling molecules, even before contact, and drive the early events of infection [8]. For telluric pathogens of the genus *Phytophthora*, these events consist sequentially of the attraction of zoospores to the root surface, followed by adhesion, aggregation and association with soil bacteria. Precedent studies indicate a key role for K^+^ gradient in zoospore aggregation in vitro [17]. Swimming Physics of zoospores has been also investigated. Through a combination of experiments and modeling, it has recently been shown how the two flagella contribute to generate the thrust when they beat together, with the anterior flagellum to which the mastigonemes are attached being the main source of thrust [35]. But to our knowledge, no detailed description of the kinetics of zoospores swimming towards a root to be infected has been made [8, 23]. Such a study could provide a better understanding of how zoospores perceive signals guiding them to the root to be infected.

Over the past decades, microfluidics has been widely used in various fields of study, in physics, biology or medicine. Microfluidics is ideal for studying cells or bacteria in controlled environments. It is particularly convenient for generating gradients to study the chemotaxis of cells or bacteria [17, 11, 33, 2]. Plant roots have also been introduced into microfluidic devices to study the development and physiology of growing roots [19] in controlled environments, or to study interactions with soil bacteria [25, 24, 5].

In this work, we use a simple microfluidic device including a living root to study the movement of zoospores swimming towards a root. We investigate the telluric species *P. parasitica*, a polyphagous pathogen attacking a wide range of hosts [29], swimming towards a root of *Arabidopsis thaliana*. Our work revealed a distance (≃300 *μ*m) from the root below which zoospores significantly change their behavior. This suggested that in our set-up, a signal is percieved at this distance from the root. Our results also show that aggregation on the root occurs at a constant rate and that this rate depends on the density of zoospores.

## Results

### The presence of the root modifies the speed distribution

To study the kinetics of zoospores approaching an *Arabidopsis* root, we built up a microfluidic device inspired by Mashala et al. [25]. It consists in a PDMS patch sticked to a glass slide (Fig. 1-A). The PDMS is molded in such a way that the root can grow into a 150 *μ*m high channel (its width is 2 or 4 mm). Typically the observations are made on a field of view containing the root apex where zoospores are expected to encyst (the distance from the apex is then no more than 2 mm) and on a field of view far from the root (at least 5 mm from the root). Using TrackMate plugin from Fiji, we reconstruct the trajectories of zoospores. We use a Matlab code to extract Fiji data, allowing us to calculate instantaneous speed.

**Figure 1.**
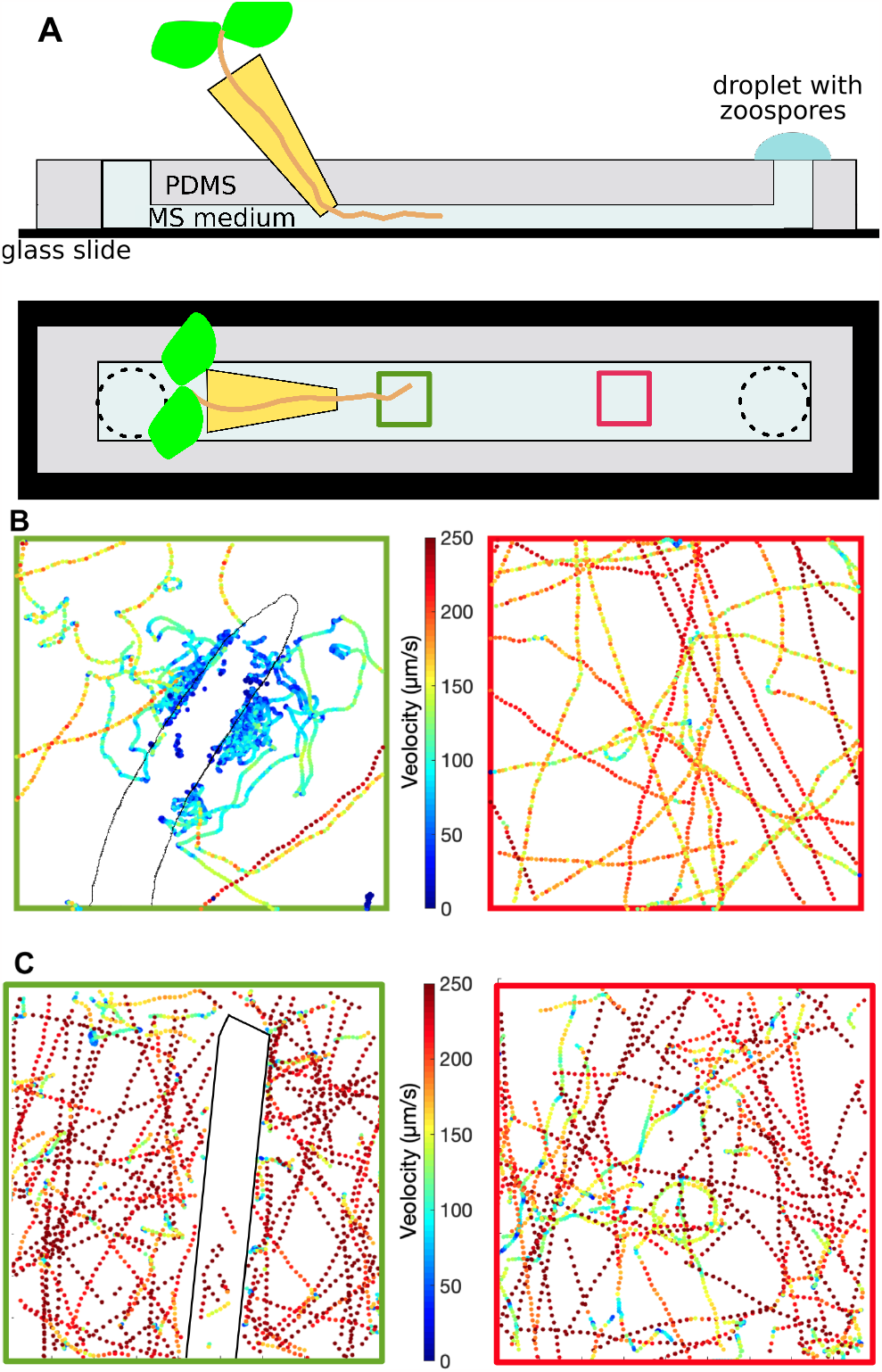
Experimental set-up and typical trajectories of zoospores close to and far from the root. (A) The sketch shows side view and top view of the microfluidic device used to grow a root in a micrometric channel and observe the zoospores swimming towards the root. A seed is allowed to grow through a pipet tip filled with agar to a microchannel filled with medium. Squares indicate fields of view (green: near the root; red: far from the root). (B) Traces represent the trajectories of zoospores tracked during 30 sec in a field of view including the root (left) and in a field of view far from the root (right). (C) Experiment realized in the same conditions but with an optical fiber instead of a root. Traces represent the trajectories of zoospores tracked during 60 sec in a field of view including the fiber (left) and in a field of view far from the fiber (right). Fields of view: 882 *μ*m x 882 *μ*m. The trajectories are colored according to the magnitude of the instantaneous speed.

Fig. 1-B shows typical trajectories observed in each field of view; The points on a trajectory are colored according to the associated instantaneous speed. Far from the root (on the right in the figure), the majority of the zoospores have straight-line trajectories with few turns and high speed; close to the root (on the left in the figure), zoospores experience a decrease in speed coupled to an increase in turns. We made five different experiments with five roots (53 movies) and all experiments showed similar zoospore behavior. We quantify these findings in the following. We also see that the zoospores swim toward and encyst on a typical site at the root tip. We then define, for each experiment, a center of aggregation where zoospores encyst.

The decrease in speed could be due to an obstacle effect due to the spatial occupation of the root. To check this point, a control experiment was performed using an optical fiber instead of a plant root. An optical fiber was chosen because it is an inert object and its size is similar to that of a root of *Arabidopsis*. In this case, zoospore trajectories do not appear to be affected by the presence of the fiber (Fig. 1-C), suggesting that the root does not act as a simple obstacle.

In order to compare the speed far from the root to the speed near the root, we can plot the instantaneous speed distribution in each field of view. Fig. 2-A shows distributions extracted from the data from Fig. 1-B. It highlights that the speed distribution is different in the presence of the root: when swimming close to the root, the speed of the zoospores significantly decreases. This is the case in all experiments and during the whole experiment (about one hour). In the control experiment the instantaneous speed distribution close to the fiber is the same as the one far from the fiber (Fig. 2-B).

**Figure 2.**
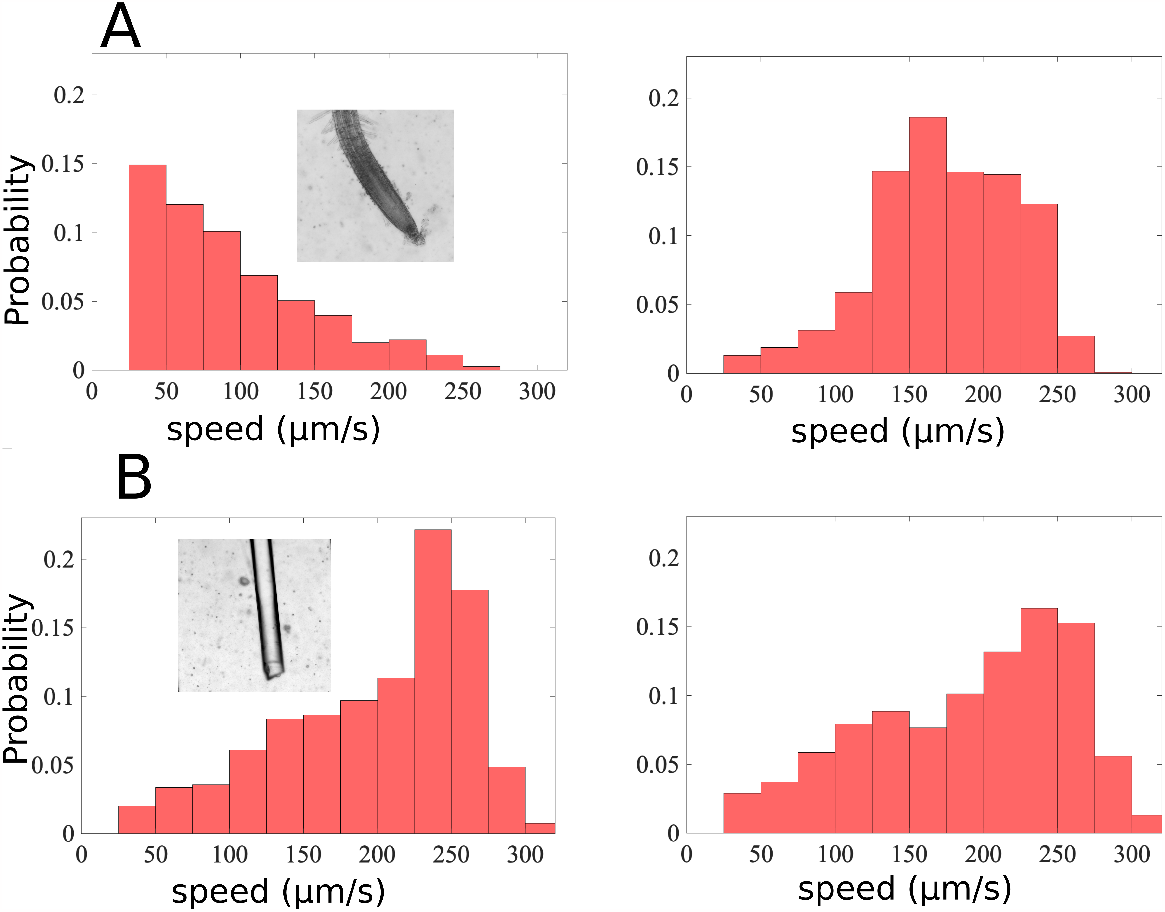
The root impacts the speed of zoospores. Instantaneous speeds are calculated: (A) in an experiment with a root introduced in the channel: in a field of view (see inset) including the root (left; 30 sec of cumulative data), and in a field of view far from the root (right; 2 minutes of cumulative data); (B) in a control experiment where the root is replaced by an optical fiber: in a field of view (see inset) including the fiber (left; 4 minutes of cumulative data), and in a field of view far from the fiber (right; 4 minutes of cumulative data). The speed distribution is modified in the presence of a root, but no significant change in speed distribution is observed in the control experiment. Insets: field of view = 882 *μ*m x 882 *μ*m.

### Speed is only impacted at short distances from the root

To go further, and find out the distance at which the speed is affected by the root, we calculate the mean speed *v*_*M*_ of zoospores as a function of the distance *d*_*agg*_ from the center of aggregation. Typical *v*_*M*_ (*d*_*agg*_) curves from one experiment are shown in Fig. 3-A, at different times during the experiment. In this experiment, the curves are shown from 16 min to 40 minutes after the start of aggregation. We do not see significant enough differences between the curves to be able to state that *v*_*M*_ (*d*_*agg*_) could be time dependent. We therefore assume that for one experiment, we can average the curves over the total duration of the experiment. Fig. 3-B presents averaged curves of three different experiments. It shows that the curves are similar from one experiment to another.

**Figure 3.**
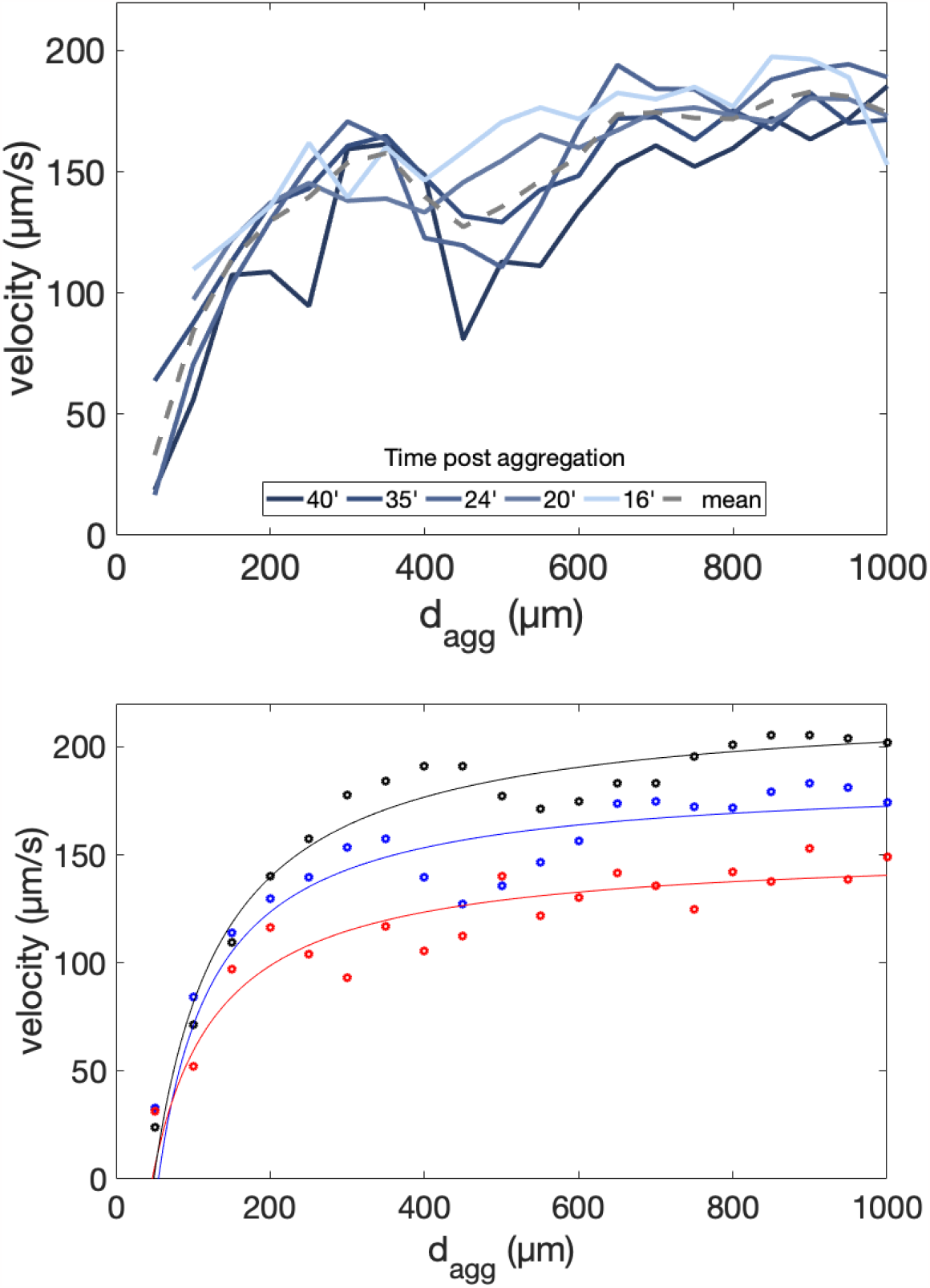
Speed as a function of the distance from the center of aggregation. (A) The speeds are calculated at different times during the same experiment (at t+16’, 20’, 24’, 35’ and 40’ after start of aggregation, from dark blue to light blue; gray dotted line: average). (B) The speeds are averaged on the total time of the experiment and plotted for three different experiments (dots). Data are fitted (lines) by a rational function *v*(*d*_*agg*_) *= c*(*d*_*agg*_ *R*_0_)/(1 + *a*.*d*_*agg*_) where *R*_0_ is the root radius at the center of aggregation, measured on the movies (see Material and methods for details)

The data were fitted (see caption) and we defined a characteristic distance *ξ* from the center of aggregation below which zoospore speed significantly decreases as 0.75 times the maximum speed (i.e. the speed far from the center of aggregation). The three experiments in Fig. 3-B give an average < *ξ > =* 295 ± 19*μm*. We shall retain that the zoospore speed is affected at a distance of about 300 *μm* from the center of aggregation.

### Directness is also impacted at short distance from the root

When zoospores swim close to the center of aggregation, they seem to experience more turns and less straight-line drives (Fig 1-B). To make it quantitative, it is possible to calculate the directness of a trajectory. When an object goes from a point *A* to a point *B*, the directness *D* is defined as the ratio between the Euclidian distance *AB* and the accumulated travelled distance between *A* and *B. D* is equal to 1 when the object moves in a straight line and tends to 0 when it makes many turns.

We evaluate directness as explained in Materials and Methods for zoospores swimming at a distance greater or less than *ξ* from the center of aggregation (Fig. 4). We sort the trajectories into curved trajectories (those that verify 0 < *D <* 0.5), and straight-line trajectories (those that verify *D >* 0.8). Fig. 4 shows that trajectories are predominantly curved at a distance of less than 300 *μ*m from the center of aggregation, in contrast to trajectories at a distance greater than 300 *μ*m. Furthermore, in the control experiment performed with an optical fiber, no change in directness is observed, emphasizing the fact that it is indeed a specific interaction between the root and the zoospores that is responsible for the change in both velocity and directness.

**Figure 4.**
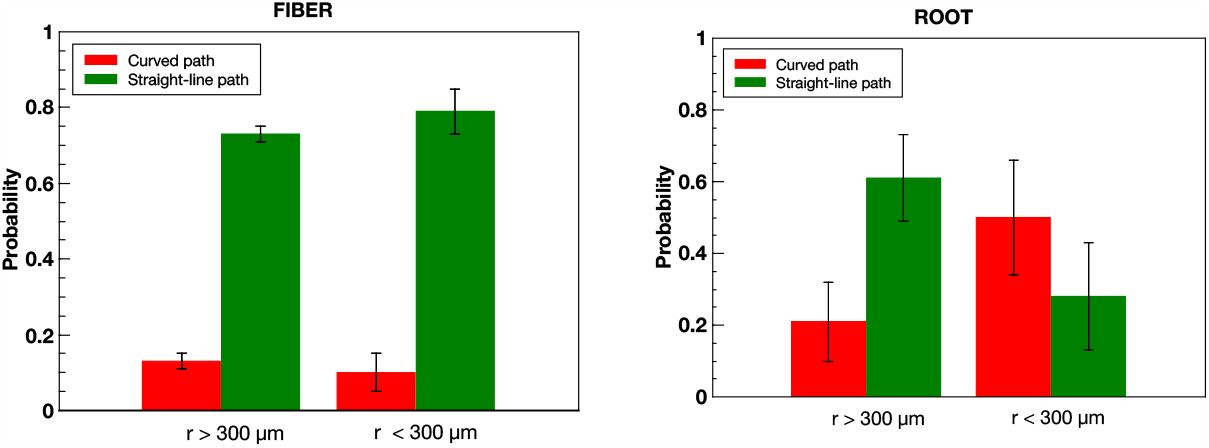
Directness is affected by the proximity of center of aggregation. Directness *D* is calculated on trajectories at a distance *r* from the center of aggregation greater or less than *ξ* (*ξ*=300 *μ*m), for the control experiment with an optical fiber (left), and for the five experiments (total of 29 movies) with a root (right) and is used to define curved trajectories (when 0 < *D <*0.5) and straight-line trajectories (when *D >*0.8). The bars show the percentage of curved trajectories in red and straight-line trajectories in green. Error bars: standard deviations.

### Zoospores aggregate on the root at a constant rate

At the very beginning of the experiment, when the density of zoospores encysted on the center of aggregation is not very high, it is possible to count the encysted zoospores. In addition, by measuring the gray intensity on the center of aggregation, it is possible to obtain an optical density (OD). By plotting OD as a function of the number of encysted zoospores, we are able to calibrate the OD measurements and use the calibration to plot an estimate of the number of encysted zoospores on the center of aggregation throughout the experiment as a function of time (Fig. 5-left).

**Figure 5.**
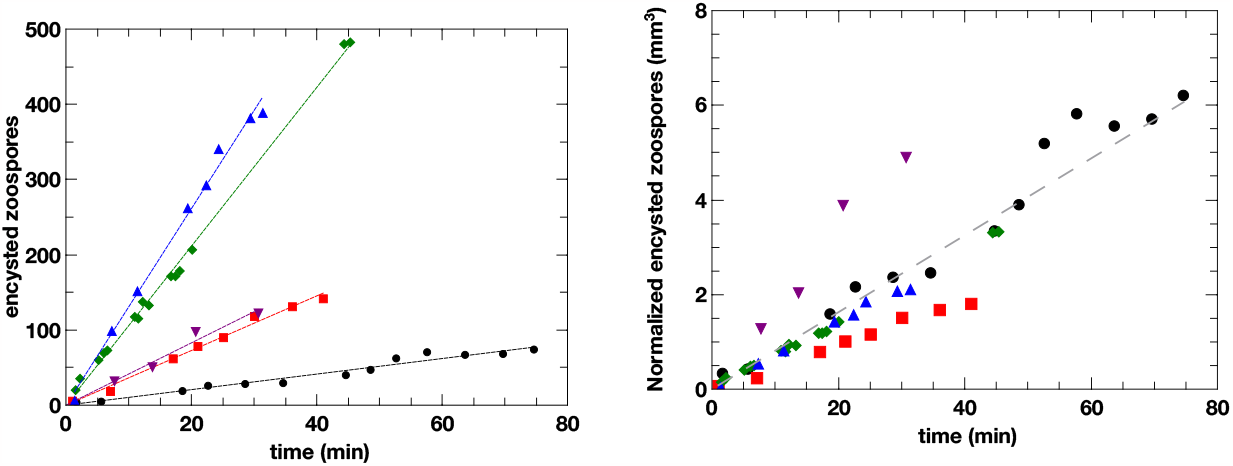
Zoospores encyst on the center of aggregation at a constant rate dependent on zoospore density. Left: The number of encysted zoospores deduced from OD measurements is plotted as a function of time for five different experiments (symbols). For each experiment, the time origin is defined as the time when the zoospores start to encyst on the center of aggregation and line is a linear fit of the data. Right: The number of encysted zoospores is normalized by zoospore density and plotted as a function of time, for all five experiments. Gray dashed line: linear fit of the data.

As a result, the number of encysted zoospores increases linearly with time for the duration of each experiment. An aggregation rate can then be calculated by fitting the data linearly. The aggregation rate varies from ≃ 1 to ≃ 14 zoospores/min. Observation of the movies shows that low (respectively high) aggregation rates correspond to experiments where the density of zoospores is low (respectively high). To be more quantitative, we evaluated the swimming zoospore density by counting the total number of zoospores in a field of view and dividing it by the volume of the field of view. The number of encysted zoospores is then normalized by the calculated density of zoospores and plotted as a function of time (Fig. 5-right). A linear fit of the normalized data gives a slope of approximately ≃ 8×10^−2^*μl* per min ≃ 5 *μl*/*h*.

## Discussion

In order to study in detail the kinetics of zoospores approaching a root, we have set up an original microfluidic device allowing the growth of an *Arabidopsis* root and the observation in real time of the swimming of zoospores moving towards the root and aggregating around the root tip. Our experiments show that the presence of the root modifies the kinetics of the zoospores: their speed greatly diminishes near the root and this decrease correlates with a decrease in their directness. A control experiment performed with an optical fiber in place of the root proved that it is not the simple physical constraints that modifies the kinetics of zoospores. A signal is then sent out by the root experienced by the zoospores within a characteristic distance *ξ* ≃ 300*μ*m from the area of aggregation.

It can be hypothesized that a signal is sent by the plant at the center of aggregation; following the detection of this signal the zoospores could increase their frequency of change of direction. It has been shown that turns are correlated with low speed [35], thus the detection of the signal could eventually lead to a decrease in the speed of zoospores. Note that we have no idea of the nature of the signal. It could be chemical or electrical [8, 23]. Further experiments will have to clarify this point.

In a laminar flow close to a surface, a velocity gradient appears near the surface, giving rise to the phenomenon of boundary layer. The thickness of the boundary layer in which the gradient is present is: 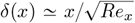 where *x* is the distance travelled on the surface and *Re*_*x*_ the local Reynolds number calculated with *x* as characteristic distance [16]. In the layer, transport of small elements is dominated by diffusion rather than advection: in this kind of region, microorganisms can then find home easier [30]. In our setup, residual flow cannot be ruled out because the pressures at the entrance and exit of the channel are not well controlled. A residual flow could then generate a boundary layer near the root, within which the chemical gradients produced by the root would not be disturbed. However, the advection velocity due to this flow does not exceed 10 *μ*m/s (data not shown). At *x* = 100*μ*m from the root tip, taking a Reynolds number of 10^−3^, a boundary layer of 3 mm thickness is expected, which is ten times larger than the characteristic distance of 300 *μ*m measured in the experiment. The 300 *μ*m distance measured cannot therefore correspond to the boundary layer due to residual flux.

What the *ξ* value of 300 *μ*m could mean for the zoospore-root interaction? Modes of colonization of root microenvironments by zoospores can vary depending on the local nature of exudates, on the biochemical composition of the mucilage and on the physical and mechanical properties of the root surface. At the soybean root cap, the Root Extracellular Trap (RET, made up of a diverse set of compounds coating border cells) prevents *P. parasitica* zoospores from colonizing the root tip early during infection [32]. Such RET was shown in various plant including *A. thaliana* [37, 14] It is now established that RET provides the first line of defense of plants against soil microbial pathogens at the root tip [13, 15, 14]. The extracellular space defined in this study by *ξ*, precisely corresponds to the microenvironment of the elongation zone at which zoospore aggregation and root infection occur [4]. Largely, this elongation zone is assumed to be the location of exudation of phytochemicals [20, 6, 7, 10] without particular subsequent architecture such as RET or mucilage [36]. In our experiments, except some AC-DCs [15] close to the tip of the root, nothing was visible close to the center of aggregation. Based on these data, the hypothesis of a slowing down of zoospore velocity resulting from the perception of a local and soluble root exudate(s) is more plausible than the assumption of a slowing down due to a physical barrier impacting zoospore movement close to the elongation zone.

We observed a constant aggregation rate (from 1 to 14 zoospores /minutes) that positively correlated with the zoospore concentration (12 to 185 cells/*μ*l). Previous work measuring the aggregation rate of *P. parasitica* zoospores at the surface of wounded leaves of *Nicotiana tabacum*, showed that 100 cells aggregated per min (zoospore suspension of 400 cells/*μ*l [18]). These results are consistent with our finding. Nevertheless, the aggregate formation rate on wounded leaves seems higher than in our conditions. This suggest that wounding generated more signals or an increase of signal intensity than intact roots. We also compared our inoculation conditions to those triggering disease development. Previous work analyzing the infection of *Arabidopsis roots* by *P. parasitica* showed that a zoospore concentration of 50-100 cells per ml was sufficient to initiate disease and invade the whole plant within 15 days [3]. In the present analysis, experiments were performed with more concentrated zoospore suspensions (100-3000 more concentrated). Thus, using here microfluidic devices with conditions that should trigger disease development if the plants could be observed later, we showed that the aggregation rate was correlated to inoculum concentration.

Our analysis of zoospore kinetics revealed a small range signal, but there could be one long-range signal whose impact on the kinetics we could not measure in our experiment. In fact, the confinement of a linear microchannel does not allow the easy testing of long range signals. However, we now discuss how the results in Fig.5 can argue for the absence of a long-range signal. Suppose that the zoospores swim isotropically at a mean speed *v*_0_. At what rate will they encounter the root tip? If we assume that the area to be infected can be assimilated to a cube of side *f* and that this cube is essentially accessible by three faces only (see Fig. 6), the number of zoospores reaching the cube during a time Δ*t* is : *N* = *v*_0_.Δ*t.ℓ*^2^.*C*/2 (considering that one sixth of the zoospores swimming isotropically go on average in a given direction of space), where *C* is the zoospore density. The aggregation rate *r* normalized by the zoospore density is : *r/C = N* (*C*Δ*t*) = *v*_0_*ℓ*^2^/2. Taking *v*_0_=150 *μ*m/s, and *ℓ*=100 *μ*m, we get *r*/*C* ≃ 3 *μ*l/h which is of the same order of magnitude as the fitted value in Fig. 5-right (5 *μ*l/h). The results therefore validate the idea of a random encounter of zoospores with the root and does not highlight the existence of a long distance signal.

**Figure 6.**
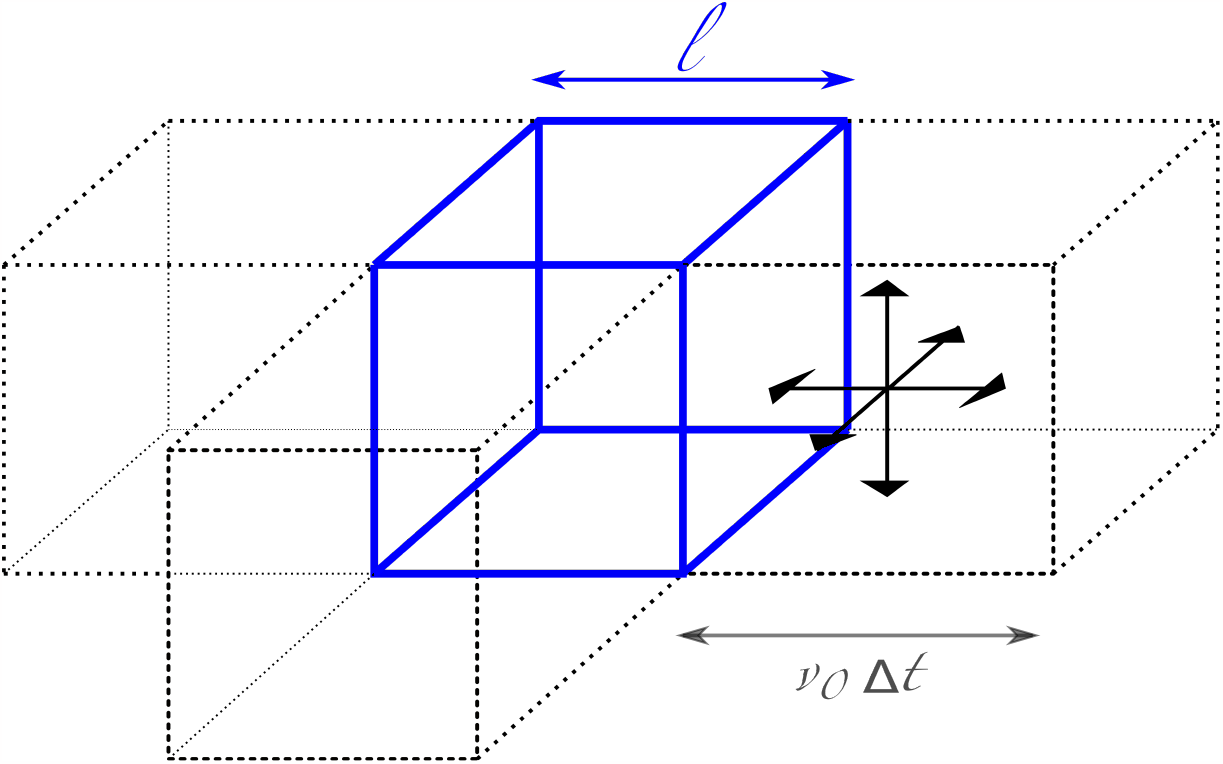
The aggregation area is assimilated to a cube of length *f*. Due to the proximity of the top and bottom surfaces, only three sides are assumed to be accessible. The density *C* of zoospores is assumed to be constant. To reach a side within Δ*t*, a zoospore swimming at speed *v*_0_ must be at a distance of at most *v*_0_Δ*t* from the side, and swim toward the side.

Past experiments suggest that aggregated zoospores themselves emit an attraction signal towards planktonic zoospores [31, 18]. Also, it is expected that the plant responds to the aggression by modifying the signals emitted at the root level. However, in our experiments, aggregation rate is constant during one hour: if the signal of attraction is regulated or if other signals are sent by the plant and/or the zoospores, none are evidenced by our experiments. Post-aggregation signals (from aggregated zoospores or from the plant) and their effect would require another device to be put in place.

## Materials and methods

### Strains and media; experimental protocol

The *Arabidopsis* ecotypes Columbia0 (Col-0), was obtained from The European Arabidopsis Stock Centre, Nottingham, UK. *Arabidopsis* seeds were surface sterilized for 5 min with sodium hypochlorite (20% commercial bleach, 2.5% active ingredient) and rinsed twice in 95% aqueous ethanol. Seeds were cold-stratified for 2 days.

*Phytophthora parasitica* Dastur isolates INRA-310 was initially isolated from tobacco in Australia, and maintained in the *Phytophthora* collection at INRA, Sophia Antipolis, France.

Murashige and Skoog (MS) medium [27] (Sigma Chemical Company) was used to cultivate plants. For seed germination, a cut pipette tip is filled with MS 0.5x supplemented by 2% sucrose in 1% agar. A seed is deposited on the agar and the tip is placed in a Petri dish filled with MS 0.5x in 2% Agar and let at 25°C. After 3-4 days, the tip is placed in the microfluidic device (see below) filled with MS 0.5x and the root is left to grow at 25°C. After another 4-5 days, the channel is rinsed with MS 0.1x before introduction of zoospores. Zoospores are recovered in water or MS 0.1x and a drop of zoospores is deposited at one inlet. Experiments are then made at room temperature.

### Microfluidic device

The microfluidic device has been fabricated using the classical soft lithography methods [26, 39]. First a mold of the channel is made by exposing a laminated film of epoxy resin (SUEX, 150 *μ*m) to UV light (wave lenght 365nm) through a plastic mask of the channel (Selba, 50600 dpi). Then the polydimethylsiloxane (PDMS, Sylgard 184, Dow corning) channel is molded by curing liquid PDMS on the resin mold at 70°C during 3h. The molded and cured PDMS channel is finally drilled with three holes (one for the inlet and outlet of solutions and one for the inlet of the growing root) and then sealed onto a glass slide by plasma treatment (Harrick plasma cleaner).

### Optical fiber

The optical fiber is a gift from Wilfried Blanc (Institut de Physique de Nice). It is a silica optical fiber obtained by drawing. Its diameter is 125 *μ*m.

### Microscopy

Observations are made under a Nikon Ti2 inversed microscope, using an objective Nikon Plan Achromat 4x (NA 0.1), and the NIS software. Images are recorded via a DFK 33UX264 (TheImagingSource). Depending on the size of field of view (from 1024 pixels x 1024 pixels to 2048 pixels x 2048 pixels), the acquisition frequency is adjusted according to the camera’s capabilities. It ranges from 8 frames per seconds to 14 fps.

### Image processing; data treatment

The acquired data were first processed with Fiji, using the TrackMate plugin. A homemade MatLab program was then used to generate velocity histograms, trajectory plots, calculation of directness and calculation of velocities as a function of distance from the aggregation zone. All fits are made using Qtiplot software.

#### Instantaneous speed

It is calculated as following:

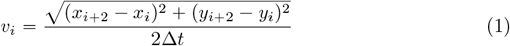

where Δ*t* is the time interval between two acquired pictures, *x*_*i*_ and *y*_*i*_ are abscissa and ordinate at instant *t*_*i*_.

#### Directness

On trajectories of at least 15 points, a directness *D* is computed for each position of the trajectory as *D* = *d*_*Euclid*_/*d*_*cumul*_ where *d*_*Euclid*_ is the Euclidian distance between the considered position and the position 15 points further and *d*_*cumul*_ is the cumulative distance travelled between the considered position and the position 15 points further.

#### Speed vs *d*_*agg*_

Speed are computed as described above. The position of center of aggregation is chosen as the middle of the region where the zoospores aggregate. When a speed is computed, the distance from center of aggregation *d*_*agg*_ is calculated as the distance between the position of the zoospore and the position of center of aggregation. For each range [*k*\*50*μ*m ; (k+1)*50 *μ*m] of *d*_*agg*_ where *k* is a positive or null integer, the computed speeds are averaged and the average is represented by a dot on the figure 3-B.

## Aknowledgements

Grants: Academy 4-Complexity and diversity of living systems, Université Côte d’Azur The authors also acknowledge the support of the French Agence Nationale de la Recherche (ANR), under grant ANR-22-CE20-0021.

